# Exploring the link between extended red blood cell parameters and platelet indices in voluntary blood donors

**DOI:** 10.1101/2024.09.29.615743

**Authors:** Ranita De, Deepak Marballi Basavaraju, Leo Stephen, Kavitha Lakshmi, Joy Mammen, Eunice Sindhuvi Edison

**Affiliations:** Department of Haematology, Christian Medical College, Vellore, India; Sree Chitra Tirunal Institute for Medical Sciences & Technology, Thiruvananthapuram, Kerala, India; Department of Transfusion Medicine and Immuno-Haematology, Christian Medical College, Vellore, India

**Keywords:** Hematology, Statistics, Laboratory Methods and Tools, Platelet Studies

## Abstract

**Background:** As regular blood donors are prone to iron deficiency, importance of extended red blood cell (eRBC) parameters in identifying donors with depleted iron stores was investigated. Thrombocytosis has been well documented in patients affected with IDA. Thus, significance of eRBC parameters in identifying iron deficiency associated thrombocytosis was also examined in this cohort.

**Methods:** Blood samples were collected in EDTA tubes from consenting donors for analyses of routine haematological and eRBC parameters. Serum samples were isolated for estimation of iron parameters.

**Results:** Iron deficient donors had significantly altered eRBC parameters. Among them, Ret-He with a cut-off of ≥32 pg had high AUC (0.822) and showed relatively high sensitivity & specificity in detecting iron deficiency. Combination of Ret-He with CCI increased sensitivity & specificity to 90.6% and 98.2%, in detection of donors affected with iron restricted erythropoiesis. This cohort had increased platelet counts, which showed significant association with Ret-He (β= -0.373), RBC-He (β= -0.384), CCI (β= 0.384), Hypo-He (β=0.494) and Micro-R (β= 0.299). Elevated platelet counts also showed significant correlations with these eRBC parameters, which was absent in iron replete donors.

**Conclusions:** eRBC parameters are sensitive indicators of non-anaemic iron deficiency, which may be enhanced by combining them. Their significant association with elevated platelet counts in iron deficient donors, highlights their importance in reflecting iron deficiency associated thrombocytosis.

**Impact Statement:** The present study discusses utility of eRBC parameters in detecting non-anaemic iron deficiency, in regular voluntary blood donors. Previous reports have investigated the importance of these parameters in identifying IDA in blood donors. The present study is the first one which indicates combining different eRBC parameters such as Ret-He and CCI increases their accuracy of detection. They were also significantly associated with elevated platelet counts in iron deficient donors, which was absent in iron replete individuals. This link between eRBC parameters and higher platelet counts in healthy donors affected with non-anaemic iron deficiency has not been reported before.

## Introduction

Voluntary blood donation is a humanitarian act towards the needy by healthy individuals. There is a need to create awareness about overall well-being and health of regular voluntary blood donors. With each donation men lose 242 ± 17 mg and women lose 217 ± 11 mg of iron (1). Although women are more susceptible to iron deficiency than men, regular blood donation is a common causative factor of iron deficiency in both sexes (2). Haemoglobin estimation by itself has poor sensitivity in the detection of early stages of iron deficiency (3). Ferritin is considered the gold standard for the diagnosis of iron-deficiency (4). However since ferritin is an acute phase reactant, any infectious or inflammatory conditions must be ruled out to prevent false positive results (5). The sensitivity of red blood cell (RBC) parameters in detecting non-anaemic iron deficiency is significantly lower. New generations of full blood count (FBC) analysers provide extended Red Blood Cell or advanced research parameters (eRBC), which have been extensively used in differential diagnosis of anaemia, thalassemia, infections and onco-haematological diseases (6). Frank Boulton *et al*. had proposed the combined cell index (CCI) as an alternative method of iron status assessment in blood donors (7).

Among the extended RBC parameters, reticulocyte Hb equivalent i.e. Ret-He facilitates early detection of iron deficiency and may serve as a valuable tool in the timely preoperative diagnosis of Iron deficiency anemia (8). The present study compares different eRBC parameters in detecting non-anaemic iron deficiency among male voluntary blood donors.

One of the common complications of IDA is thrombocytosis. While most patients affected with IDA have normal platelet counts, variable numbers of affected individuals develop thrombocytosis (9,10). Some patients suffering from severe IDA develop thrombocytopenia, which normalized after iron therapy (11).We investigated the association of iron parameters and eRBC parameters with elevated platelet counts, observed in iron deficient donors. Univariable linear regression and subsequent multivariable linear regression was utilized to understand the significance of haematological parameters, in identifying thrombocytosis associated with non-anaemic iron deficiency.

## Materials and methods

### 1. Settings

The prospective observational study was approved by institutional review board and was conducted from November 2017 to May 2021 in the Departments of Haematology and Transfusion Medicine and Immuno-Haematology at Christian Medical College, Vellore, India. Donors were initially subjected for screening in accordance with Directorate general of Health Services (DGHS), India guidelines (12).

## 2. Subjects

Consenting male voluntary blood donors aged between 18-55 years with Hb ≥ 12.5 g/dl were included in the study. Female donors were not included as they are more prone to iron depletion due to physiological blood loss. Donors who were not consenting and/or on iron/vitamin supplementations were excluded from the study.

Pre-donation haemoglobin screening was by copper sulphate method and confirmed by automated haematology analyser counter. The subjects who fulfilled the inclusion criteria were administered with a short questionnaire to collect data on frequency of donation in last 2 years, body weight and height [body mass index (BMI)], smoking habits, alcohol, and meat consumption. Donors were categorized as low frequency (1-2 donations in last 2 years), moderate frequency (3-4 donations in last 2 years) and high frequency (≥ 5 donations in last 2 years)(7). Smoking index was taken into consideration to classify donors as mild, moderate and heavy smokers (13).

## 3. Analytical methods

Samples collected in EDTA tubes were processed within 4 hours for analysis of various haematological parameters. Apart from routine haematological parameters, eRBC parameters were measured using Sysmex XN-9000 (Sysmex Corporation, Kobe, Japan) haematology analyser. Daily quality control was performed prior to patient analysis.

Serum samples isolated from peripheral blood of donors were stored at -80^0^ C for further analyses of iron related parameters, following testing for transfusion transmissible infections (TTI). Serum ferritin levels were determined on Siemens ADVIA Centaur analyser by two-site sandwich chemiluminescence immunoassay. sTfR levels were determined on Roche Cobas 8000 modular analyser using Tina-quant Roche kit by Immunoturbidimetric method. The transferrin receptor-ferritin index (sTfR-F index) was defined as the ratio of soluble transferrin receptor level to log ferritin level (14). Serum hepcidin levels were estimated by ELISA method (Hepcidin 25 bioactive ELISA, DRG International, Inc.). Based on serum ferritin levels, donors were classified as having absent iron Stores (ferritin < 15 ng/ml) (15) and iron replete individuals (ferritin ≥ 15 ng/ml).

## 4. Statistical methods

Quantitative variables were reported using Mean ± Standard deviation (SD) or Median and interquartile range (IQR), according to the characteristics of the distribution of the data. For qualitative variables, number and percentages were used.

Comparison of different haematological parameters between donors with depleted iron stores (Ferritin < 15 ng/ml) (n=51) and iron replete donors (Ferritin ≥ 15 ng/ml) (n=241) was done by ANOVA (Graphpad Prism 8.0.1). The latter was also used for comparison of different haematological parameters between donors with normal BMI (18.5-24.9) and overweight (BMI: 25-29.9) and obese (BMI: ≥30) donors. Any association(s) between continuous variables were estimated by Pearson’s correlation coefficient. Individual associations of routine haematological and eRBC parameters with platelet counts were examined by regression analyses (IBM SPSS Statistics, version 21.0) in iron deficient donors and controls.

Receiver operating characteristic (ROC) analysis with Area Under the Curve (AUC) calculation, was employed to compare the diagnostic efficiency of eRBC parameters in detecting iron depletion. The Youden index was used to identify optimal cut-off values of these parameters for identifying donors with absent iron stores.

### 4.1. Combination of eRBC parameters to detect absent iron stores in donors with greater accuracy

Ret-He has gained prominence as an iron status marker as it provides an indirect measurement of the functional iron available for production of reticulocytes. The Reticulocyte Hb equivalent (Ret-He) is a log transformation of Ret-Y, which is the mean value of the forward-scattered-light histogram of the reticulocyte population, analysed by flow cytometry (16). An alternative method for iron status assessment can also be done by calculation of the Combined Cell Index (CCI). It is estimated according to the formula: RDW × 10^4^ × MCV^-1^ × MCH^-1^ (7).

Different eRBC parameters were considered and the resultant sensitivity and specificity after combining them were calculated using the formulae: 1 – (1 – sensitivity of test 1) × (1 – sensitivity of test 2) and 1 – (1 – specificity of test 1) × (1 – specificity of test 2) respectively (17). For all tests, p-value of < 0.05 was considered as being statistically significant. All data were analysed using Stata version 15.0 software.

## 5. Results

### 5.1. Characteristics of study cohort

Two hundred and ninety-two, consenting male voluntary donors were enrolled in the study. The median age of the study subjects was 29 years (18–55). Most of the subjects were low frequency donors (n=108, 37%) or moderate donors (n=103, 35.2%) while individuals donating ≥ 5 times in the last 2 yrs. were 14.4% (n=42).

A general trend towards obesity was apparent as about 69.9% (n=204) of the donors were overweight (BMI: 25-29.9) or obese (BMI: ≥30). Majority of the donors were non-smokers, 84.9% (n=248) and 77.4% (n=226) did not consume alcohol. Most of the subjects consumed a meat-based diet, 93.8% (n=274) (Supplemental Table 1). While 17.6% (n=51) donors had absent iron stores, majority of them i.e. 82.5% (n=241) were iron replete (Ferritin ≥15 ng/ml). Iron status of donors is summarized in Supplemental Figure 1.

### 5.2. Haematological and iron related parameters of donors with depleted iron stores

Haematological and iron parameters are presented in Table 1 (7,18,19,20,21,22,23,24). Iron stores were significantly decreased in iron-deficient donors (ferritin: 10.2 ng/ml (4.6-14.6 ng/ml) compared with iron replete individuals (ferritin: 61.81 ng/ml (15.1-658.1 ng/ml) (p< 0.001). Thus, the former cohort had significantly lower hepcidin levels (0.82 ng/ml, 0.16-34.7 ng/ml) compared to the remaining donors (8.9 ng/ml, 0.11-70.7 ng/ml) (p= 0.009). Other iron related parameters including sTfR, sTfR-F index was significantly higher (p< 0.001).

**Table 1:**
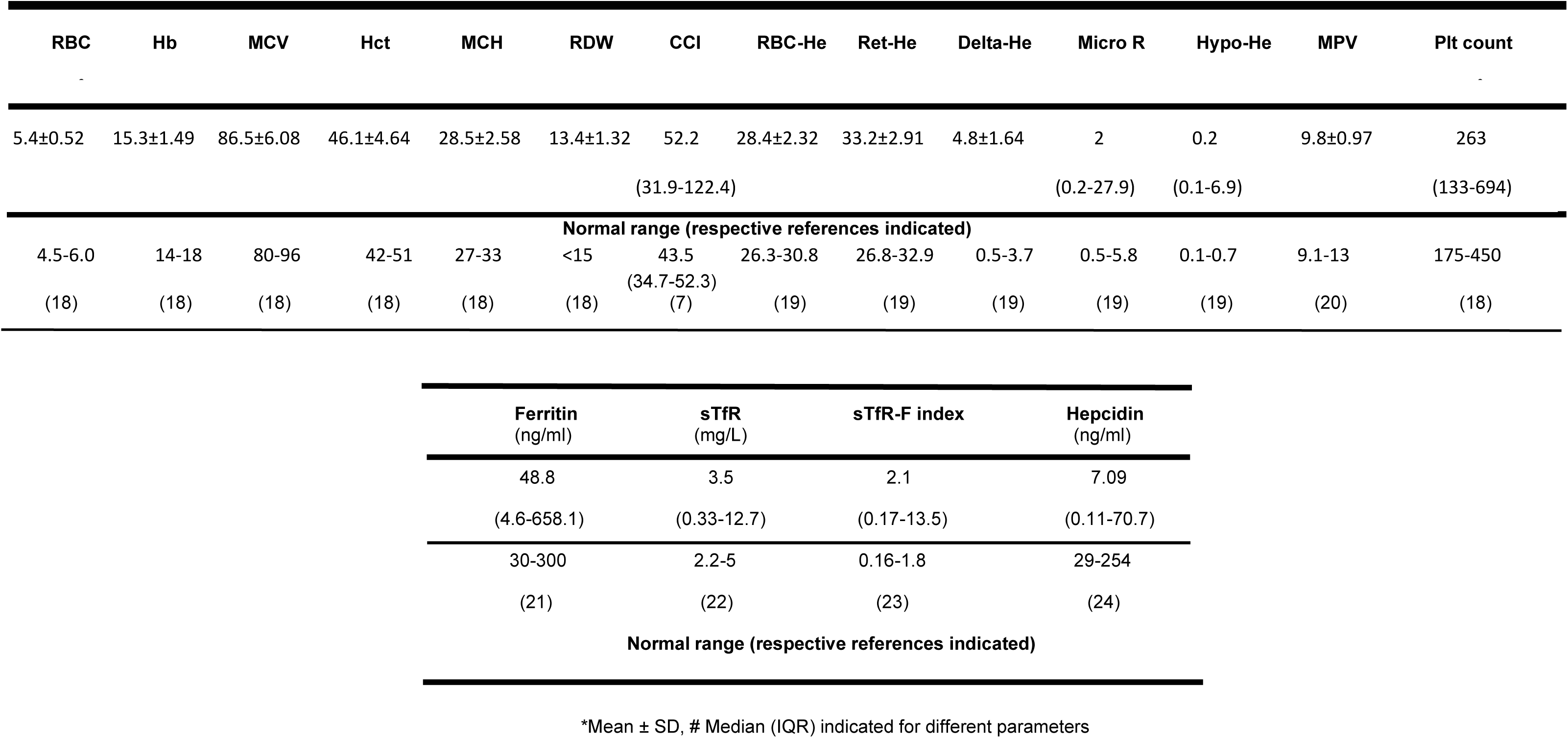
Haematological and iron parameters of voluntary donors (n = 292)

The mean Hb (14 ± 1.24 g/dL) in iron deficient donors was in the normal reported range for healthy adult males (25), indicating absence of anaemia. However, iron deficiency may have attributed to iron restricted erythropoiesis in these donors. This was indicated by significantly decreased red blood cell indices including Hb, MCV, MCH, RDW (p< 0.001) compared to controls. Reticulocyte Hb content i.e. Ret-He was also significantly lower (p< 0.001), compared to controls. Other eRBC parameters such as CCI, MicroR (% of RBCs with MCV < 60fL) and Hypo-He (% of RBCs with MCH < 17pg) were significantly higher (p<0.001) in this cohort.

The platelet counts were also affected by absent iron stores. Closely reflecting IDA associated thrombocytosis, iron deficient donors had higher platelet counts (277 × 10^3^/μl, 135-694 × 10^3^/μl), than iron replete donors (259 × 10^3^/μl, 133-435 × 10^3^/μl). This difference was not statistically significant (Table 2).

**Table 2:** Comparison of haematological and iron parameters between iron-deficient donors and controls.

| Parameter | Donors with<br>depleted iron<br>stores<br>(ferritin <15ng/ml) | Donors with<br>ferritin ≥<br>15ng/ml | p value |
| --- | --- | --- | --- |
|  | n=51 | n=241 |  |
| *Hb (g/dL) | 14 ± 1.24 | 15.5 ± 1.42 | < 0.001 |
| *MCV (fl) | 82.1 ± 6.4 | 87.4 ± 5.6 | < 0.001 |
| *MCH (pg) | 25.9 ± 2.5 | 29.1 ± 2.2 | < 0.001 |
| *RDW (%) | 14.6 ± 1.5 | 13.1 ± 1.1 | < 0.001 |
| *RBC-He (pg) | 25.9 ± 2.5 | 28.9 ± 1.9 | < 0.001 |
| *Ret-He (pg) | 30.3 ± 2.9 | 33.8 ± 2.5 | < 0.001 |
| *CCI | 70.9 ± 1.7 | 52.7 ± 9.6 | < 0.001 |
| #Micro-R (%) | 6.4 (0.3-27.9) | 1.8 (0.2-16.4) | < 0.001 |
| #Hypo-He (%) | 1.3 (0.1-6.9) | 0.2 (0.1-4.6) | < 0.001 |
| #Plt Count<br>(×10<sup>3</sup>/μl) | 277 (135-694) | 259 (133-435) | 0.97 |
| #Ferritin (ng/ml) | 10.2 (4.6-14.6) | 61.8 (15.1-658.1) | < 0.001 |
| #sTfR (mg/L) | 5.2 (1.2-12.7) | 3.3 (0.3-12.1) | < 0.001 |
| #sTfR-F index | 5.2 (1.8-13.5) | 1.9 (0.17-5.7) | < 0.001 |
| #Hepcidin<br>(ng/ml) | 0.82 (0.16-34.7) | 8.9 (0.11-70.7) | 0.009 |

\*Mean ± SD, # Median (IQR) indicated for different parameters

### 5.3. Utility of eRBC parameters in identification of non-anaemic iron deficiency

eRBC parameters have been utilized to detect IDA in various donor groups (7,8,9). The significance of these parameters in identifying iron deficiency, not leading to anaemia in voluntary blood donors, were compared by ROC curves.

Different eRBC parameters including Ret-He (AUC: 0.822, 72.6% sensitivity, 77.8% specificity), CCI (AUC: 0.825, 66% sensitivity, 91.9% specificity), RBC-He (RBC Hb equivalent) (AUC: 0.818, 68.6% sensitivity, 87.1% specificity) and Hypo-He (AUC: 0.851, 60.8% sensitivity, 95.6% specificity) displayed good diagnostic accuracy in predicting absent iron stores. Comparison of these parameters indicated that Ret-He with a cut-off of ≥ 32 pg had a high AUC (0.822) and relatively greater sensitivity and specificity, thus emerging as the best indicator of iron deficiency (p < 0.001). However, combination of Ret-He with another eRBC parameter displaying good AUC i.e. CCI, improved diagnostic accuracy. Both sensitivity as well as specificity, significantly increased to 90.6% and 98.2%, respectively. ROC curves comparing different eRBC parameters have been depicted in Figure 1.

**Figure 1.**
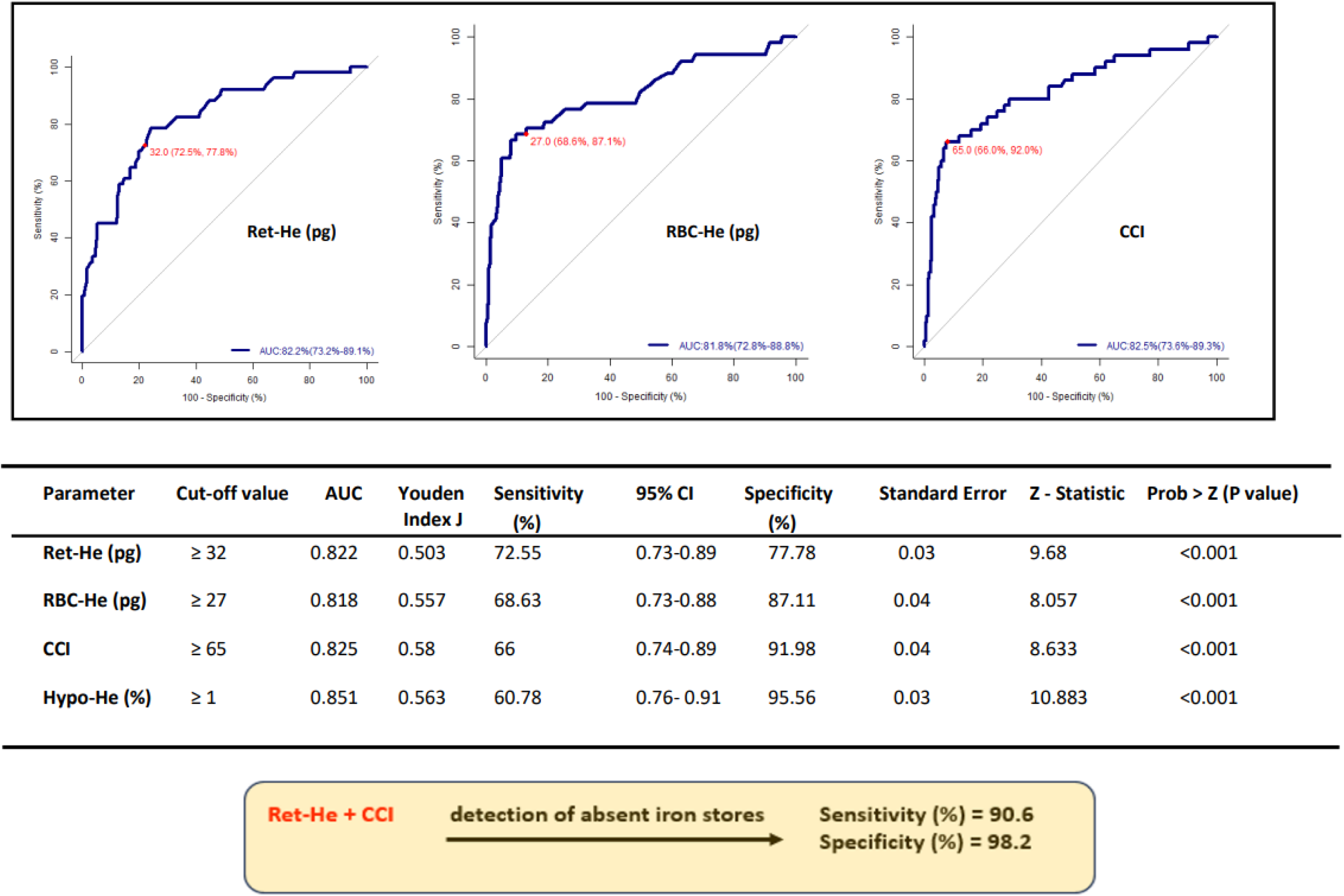
Comparison of receiver operating characteristic (ROC) curve analyses of eRBC parameters for identification of donors with depleted iron stores (ferritin < 15 ng/ml).

### 5.4. Effect of blood donation frequency on haematological and iron related parameters

As regular blood donation has been reported to adversely affect some haematological parameters (26), we investigated the association of donation frequency with routine haematological, eRBC and iron related parameters. Apart from Hb, MCV, MCH, MCHC, several eRBC parameters including RBC-He (r=-0.389, p=0.001), Ret-He (r=-0.296, p=0.001) also significantly decreased as frequency of blood donation increased. While other eRBC parameters such as CCI (r=0.402, p=0.001), MicroR (r=0.343, p=0.001) and Hypo-He (r=0.276, p=0.001) showed significant positive correlation with donation frequency (Supplemental Figure 2).

Iron stores were adversely affected by donation frequency as indicated by significant negative correlation, observed between frequency of donation and serum ferritin (r=-0.223, p=0.001) and log Ferritin (r=-0.347, p=0.001). sTfR-F index which has been reported to increase in iron deficiency (27), showed significant positive correlation with donation frequency (r=0.249, p=0.001) Thus, a relatively high percentage of moderate and high frequency donors (49.6%) may explain the presence of depleted iron stores in 17.6% of regular voluntary donors.

### 5.5. Exploring the effect of donor BMI on body iron status and platelet counts

Dietary preferences and present sedentary lifestyles often tip the BMI scale towards obesity. Most of the donors (93.8%) preferred a meat-based diet and a majority (69.9%) of them were overweight/obese. We also observed that a high percentage of donors with ferritin content < 30ng/ml were overweight/obese (62.8%).

They had higher hepcidin levels (median=2.99 ng/ml; 0.32-70.6 ng/ml), compared to donors having normal BMI (median=1.19 ng/ml; 0.11-34.7 ng/ml). The difference was not significant. The probable presence of low-grade subclinical inflammation commonly associated with obesity, may be responsible for elevated hepcidin levels.

An interesting observation was occurrence of significant positive correlation between hepcidin levels and platelet counts (r= 0.594, p < 0.001) in donors with absent iron stores (ferritin < 15ng/ml). Similar results were observed in donors with ferritin < 20 ng/ml (r=0.509, p < 0.001) and those with ferritin < 30 ng/ml (r=0.383, p < 0.001). The significance of this association was not clear, and may be investigated further in larger cohorts.

### 5.6. Exploring the link between eRBC parameters and altered platelet indices in iron deficient donors

#### 5.6.1. Correlation between eRBC parameters and platelet indices

Thrombocytosis is a common occurrence in patients affected with mild to moderate IDA (9) and certain eRBC parameters have also been reported to be elevated in the latter (28). We observed eRBC parameters were significantly altered in iron deficient donors, apart from enabling their precise identification. As platelet counts were elevated in this cohort, we compared the potential association of haematological parameters and platelet indices in iron deficient and iron replete donors.

Although iron deficient donors had significantly altered iron parameters compared to controls, they did not show any significant association(s) with platelet indices. However, a decline in MPV in iron deficient donors, was associated with an increase in CCI, MicroR and Hypo-He, indicated by significant negative correlations (CCI (r=-0.435, p=0.002), MicroR (r=-0.408, p=0.003), Hypo-HE (r=-0.351, p=0.012). Past studies have shown a negative association between MPV and platelet counts, in IDA patients (29). Similarly, we also observed that MPV showed significant negative correlation with platelet counts (r=-0.528, p < 0.001) in the iron deficient cohort (Supplemental Figure 3). No significant associations between MPV and eRBC parameters were observed in iron replete donors.

Further indication of a link between iron deficiency and elevated platelet counts was evident from significant positive correlations between platelet counts and eRBC parameters. We found CCI (r=0.380, p=0.007), MicroR (r=0.302, p=0.031) and Hypo-HE (r=0.496, p < 0.001) were positively correlated with platelet counts in iron deficient (Figure 2). These associations were absent in controls.

**Figure 2.**
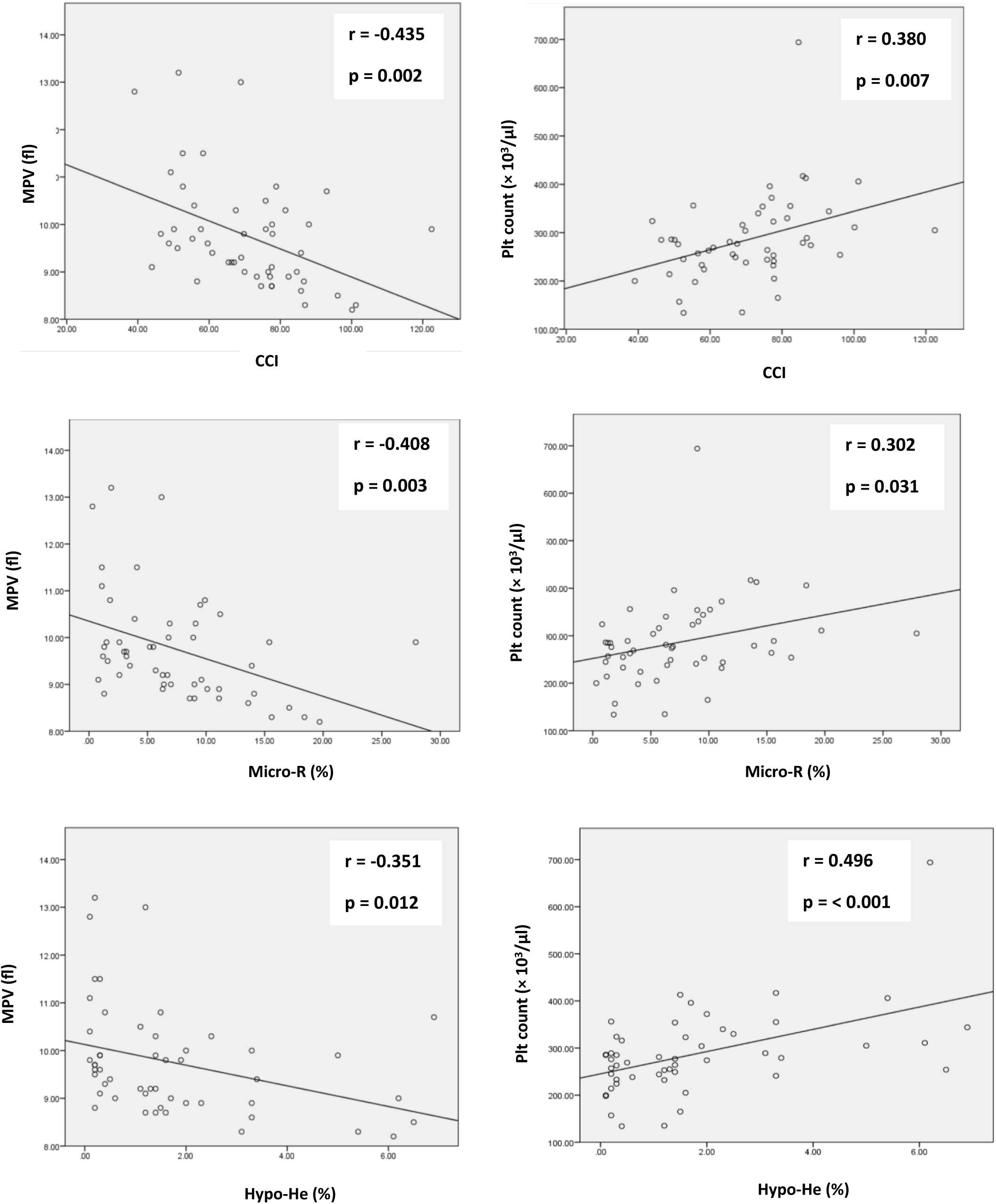
Correlation between eRBC parameters and platelet indices in donors with depleted iron stores.

#### 5.6.2. Investigating how change in haematological parameters influence platelet counts

The association between change in platelet counts and hematological parameters was further confirmed by univariable linear regression, adjusted for age, BMI, and donation frequency. While we found significant negative linear associations of platelet counts with Hb (β= -0.401, p=0.004), MCV (β= -0.286, p=0.042) and MCH (β= -0.349, p=0.012) in iron deficient donors, no significant associations were observed in controls. Some eRBC parameters including Ret-He (β= -0.373, p=0.007) and RBC-He (β=-0.384, p=0.005) showed significant negative association with platelet counts. Others such as CCI (β= 0.384, p=0.006), Micro-R (β= 0.299, p=0.033) and Hypo-He (β= 0.494, p<0.001) displayed significant positive associations with the former. In controls, however eRBC parameters did not show any significant association with platelet counts. Thus, iron deficiency adversely affects red blood cell indices and reticulocyte production. eRBC parameters including CCI, Hypo-He and Micro-R increase during iron deficiency, alongside platelet counts. Associations between hematological parameters and platelet counts have been summarized in Table 3.

**Table 3:** Association of haematological parameters and platelet counts in iron-deficient donors and controls.

| Haematological<br>Parameters<sup>b</sup> | Platelet counts<sup>a</sup><br>in iron deficient donors (Ferritin < 15ng/ml) |  |  | Platelet counts<sup>a</sup><br>in controls (Ferritin ≥ 15 ng/ml) |  |  |
| --- | --- | --- | --- | --- | --- | --- |
|  | β Coef | p-value | Adjusted<br>R<sup>2</sup> | β Coef | p-value | Adjusted<br>R<sup>2</sup> |
| Hb (g/dL) | -0.401 | 0.004 | 0.143 | -0.085 | 0.189 | 0.003 |
| MCV (fl) | -0.286 | 0.042 | 0.063 | -0.058 | 0.371 | 0.001 |
| MCH (pg) | -0.349 | 0.012 | 0.104 | -0.035 | 0.584 | -0.003 |
| RDW (%) | 0.370 | 0.007 | 0.120 | 0.077 | 0.255 | 0.002 |
| Ret-He (pg) | -0.373 | 0.007 | 0.121 | -0.069 | 0.284 | 0.001 |
| RBC-He (pg) | -0.384 | 0.005 | 0.130 | -0.101 | 0.129 | 0.006 |
| CCI | 0.384 | 0.006 | 0.130 | 0.084 | 0.198 | 0.003 |
| Micro-R (%) | 0.299 | 0.033 | 0.071 | 0.079 | 0.235 | 0.002 |
| Hypo-He (%) | 0.494 | < 0.001 | 0.229 | 0.071 | 0.290 | 0.001 |

\*a- Dependent variable – Platelet count ( $\times 10^3/\mu\text{l}$ ), b- Predictors

Upon stratification by significant interacting variables, the magnitudes of the associations between most of the hematological parameters and platelet counts were not significant. However, the negative linear association of Hb with platelet counts (β= -0.40, p=0.003) was maintained in iron deficient donors (Figure 3).

**Figure 3.**
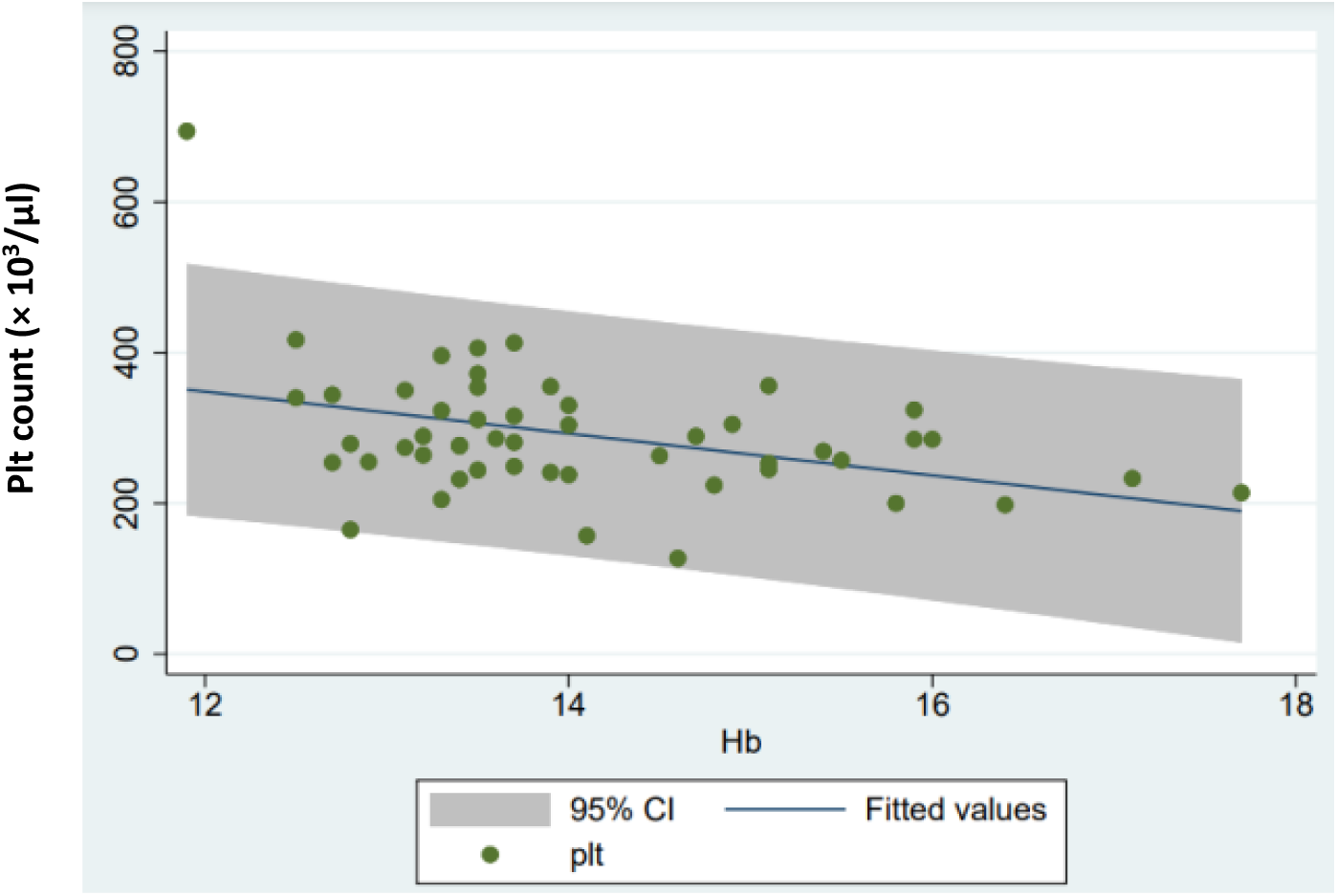
Relationship between Hb and platelet counts in iron deficient donors.

## 6. Discussion

Iron deficiency is a common complication among voluntary donors and less attention is given for their health status. Majority of the subjects in the present study were low frequency or moderate donors and were either overweight or obese, which was similar to other studies (26) eRBC parameters have gained prominence in the past decade, for sensitive detection of IDA among blood donors (26,30).

We observed that donors with absent iron stores (Ferritin < 15 ng/ml) had higher value of mean Ret-He, than those presenting with iron deficiency, in a study by Mehmet Ali Uçar *et al*. Inclusion of only male voluntary donors as well as use of a different cut off value of ferritin for defining iron deficiency, may account for this difference. However, iron deficiency was linked with decreased Ret-He values in both cases (31). Certain other eRBC parameters such as Hypo-He and MicroR were significantly higher in iron deficient donors, compared to controls. Similar trends were observed by Angeli Ambayya *et al.* in women (32). The reference interval of another eRBC parameter which increases during iron deficiency i.e. CCI was higher in our study, compared to that reported by Vuk *et al*. This may be due to a lower cut off value of ferritin used for defining iron depletion, as well as a larger cohort size in the latter study (7).

Median ferritin levels in our study was significantly lower than that observed by Reddy K V *et al.* in male voluntary donors (33). We also noted that prevalence of donors having depleted iron stores was lesser, than that recently reported by Patrick Adu *et al.* (34). One of the underlying reasons may be different ethnic backgrounds and dietary habits of the investigated cohorts.

As donation frequency increased several routine haematological parameters including Hb, MCH, MCHC decreased. Other groups have reported similar findings (8,26,30). Donation frequency influenced eRBC parameters as well, and CCI, MicroR and Hypo-He increased, with increase in frequency of blood donation. Iron stores were also adversely affected by frequency of blood donation, indicated by significant decline in serum ferritin and log ferritin, with increased donations. Similar observations have been reported before (8,26).We observed that as donation frequency increased, sTfR-F index also rises. As the latter increases in iron deficiency (27), this finding further validates the adverse effect of frequent blood donation on donor iron stores. The present study is the first one, to the best of our knowledge, which investigates the effects of donation frequency on eRBC parameters. Significant increase in CCI, Micro-R and Hypo-He in response to donation frequency, also emphasizes the importance of these parameters as sensitive indicators of iron deficiency.

In the present study, more than 60% of donors with low ferritin values (< 30 ng/ml) were overweight/obese, which may result in the presence of low-grade subclinical inflammation. This in turn may induce hepcidin production, which inhibits iron absorption and decreases serum iron levels by its sequestration in the reticuloendothelial system (35).Consequently ferritin levels have been reported to be higher in overweight/obese people (36). However, an earlier study by Eftekhari *et al.* found that ferritin levels were lower in obese adolescents (37). Thus, hepcidin levels may be elevated in donors with low iron stores, as reported by us.

We compared different eRBC parameters, to identify donors with depleted iron stores. Our results indicated that Ret-He with a cut-off of ≥ 32 pg had a high AUC (0.822) and relatively high sensitivity and specificity, emerging as the best candidate. This value was similar to an earlier report by Ucar *et al* where RET-He with a cut-off of < 35.5pg predicted iron depletion with 100% sensitivity and 55.6% specificity. They predicted a high diagnostic value of Ret-He (AUC: 0.881) in predicting iron deficiency in blood donors (31). Several studies have reported a wide range of diagnostic cut-off values of Ret-He, ranging from 25-35 pg with varying degrees of sensitivity and specificity, for assessing IDA in different study populations (38,39).

The detection sensitivity of haematological parameters may be increased by combining different haematological indices, as done by CCI. This shows high diagnostic value in predicting absent iron stores in donors (AUC: 0.961) as reported by Vuk T. *et al*. (7) We also observed that CCI (AUC: 0.825) may be a good indicator of predicting iron deficiency. While most studies have utilized eRBC parameters to detect IDA, we ascertained their significance in identifying non-anaemic iron deficiency. The present study is the first one, to show that combining two different eRBC parameters such as Ret-He and CCI, can significantly improve both sensitivity and specificity of detection.

Thrombocytosis has been reported to be a common occurrence in patients affected with IDA (9). Different research groups have investigated possible mechanisms by which iron may influence platelet biogenesis through *in vitro* and *in vivo* studies (40). In the present study, we did not find any significant correlation between iron parameters and platelet counts in iron deficient donors. Thus, we investigated if eRBC parameters displayed any association with platelet indices, in presence of iron deficiency.

We found that CCI, Hypo-He and MicroR showed significant negative correlation with MPV, and were positively correlated with platelet counts in iron deficient donors. No such associations were observed in iron replete donors. Past studies have indicated MPV to be negatively associated with platelet counts (29), which was in agreement with our findings. These results indicated a close link between iron deficiency and elevated platelet counts.

Regression analyses indicated that increase in platelet counts was significantly associated with decline in routine haematological parameters, in iron deficient donors. A similar trend was observed in eRBC parameters. While Ret-He and RBC-He showed significant negative association, CCI, Hypo-He and Micro-R increased in iron deficiency and were positively associated with platelet counts. We did not find any significant association between change in eRBC parameters and platelet counts in iron replete donors. The present study is the first one, to the best of our knowledge which investigates the link between eRBC parameters and platelet counts in voluntary donors, affected with iron deficiency.

## 7. Conclusions

eRBC parameters are sensitive indicators of iron restricted erythropoiesis in voluntary blood donors, with depleted iron stores. Their sensitivity and specificity significantly increased upon combination of different parameters such as Ret-He and CCI, which individually provide good diagnostic efficacy in detecting iron deficient donors. This may be especially useful in identification of non-anaemic iron deficiency, when an individual eRBC parameter may not provide adequate precision. These parameters also showed significant association with elevated platelet counts in iron deficient donors. Thus, a novel role of eRBC parameters in identification of iron deficiency associated thrombocytosis emerges, which necessitates further validation in IDA patients.

## Acknowledgements

RD performed experiments and analyses, wrote, edited the manuscript. Deepak MB performed experiments, edited the manuscript. LS helped in sample collection and processing. KL helped in statistical analyses. JM participated in research design, data analyses, and reviewed the manuscript. ES designed the research proposal, analysed data, reviewed the manuscript. All authors read and approved the final version of the manuscript.

## List of abbreviations

RBC parameters: Red blood cell parameters
FBC Analysers: Full blood count analysers
eRBC parameters: Extended red blood cell parameters
CCI: Combined cell index
Ret He: Reticulocyte Hb equivalent
IDA: Iron deficiency anaemia
DGHS: Directorate general of Health Services India
BMI: Body mass index
TTI: Transfusion transmissible infections
sTfR-F index: Transferrin receptor-ferritin index
ROC: Receiver operating characteristic analyses
AUC: Area under the curve
Micro-R** Hypo-He**, RBC-He** (**: Expanded in the manuscript).

