## Supplemental table 1 and Figures for "Exploring the link between extended red blood cell parameters and platelet indices in voluntary blood donors"

### Supplemental Table 1- General characteristics of voluntary blood donors (n=292)

| Age (yrs) | Frequency of donation (last 2 yrs) | | | | Smoking (Number of cigarettes/week) | | | | Alcohol consumption | | | | Body mass index (BMI) | | | | Diet | |
|---|---|---|---|---|---|---|---|---|---|---|---|---|---|---|---|---|---|---|
| 29 (18-55) | **Nil** | **Low (1-2)** | **Intermediate (3-4)** | **High (≥5)** | **Nil** | **Mild (1-3)** | **Moderate (4-6)** | **Heavy (7-10)** | **Nil** | **Occasional** | **Light** | **Heavy** | **Under weight(< 18.5)** | **Normal (18.5-24.9** | **Over weight (25-29.9)** | **Obese (≥30)** | **Veg** | **Meat-based** |
| **N** | 39 | 108 | 103 | 42 | 248 | 38 | 4 | 2 | 226 | 57 | 7 | 2 | 8 | 80 | 160 | 44 | 18 | 274 |
| **%** | 13.4 | 37 | 35.3 | 14.4 | 84.9 | 13 | 1.4 | 0.7 | 77.4 | 19.5 | 2.4 | 0.7 | 2.7 | 27.4 | 54.8 | 15.1 | 6.2 | 93.8 |

**Supplemental Figure 1- Iron status of voluntary blood donors (n=292)**

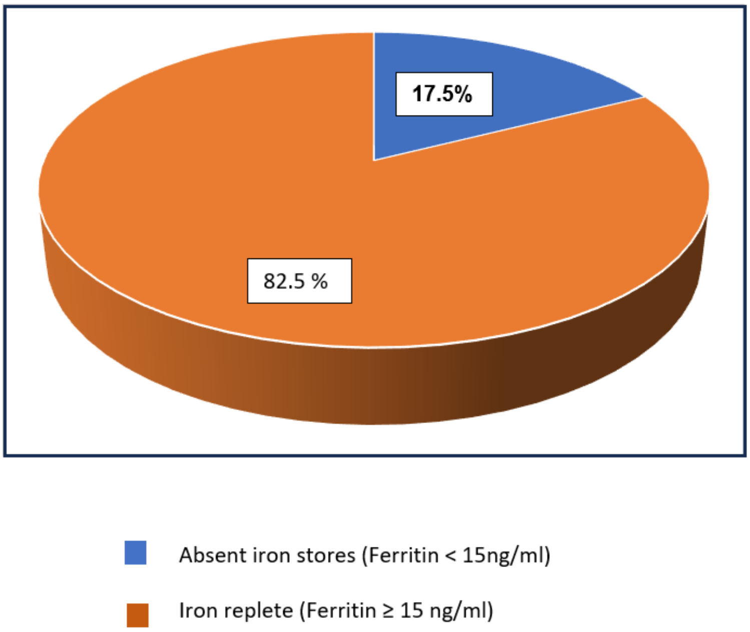

**Supplemental Figure 2- Effect of blood donation frequency on eRBC parameters and iron parameters in voluntary donors**

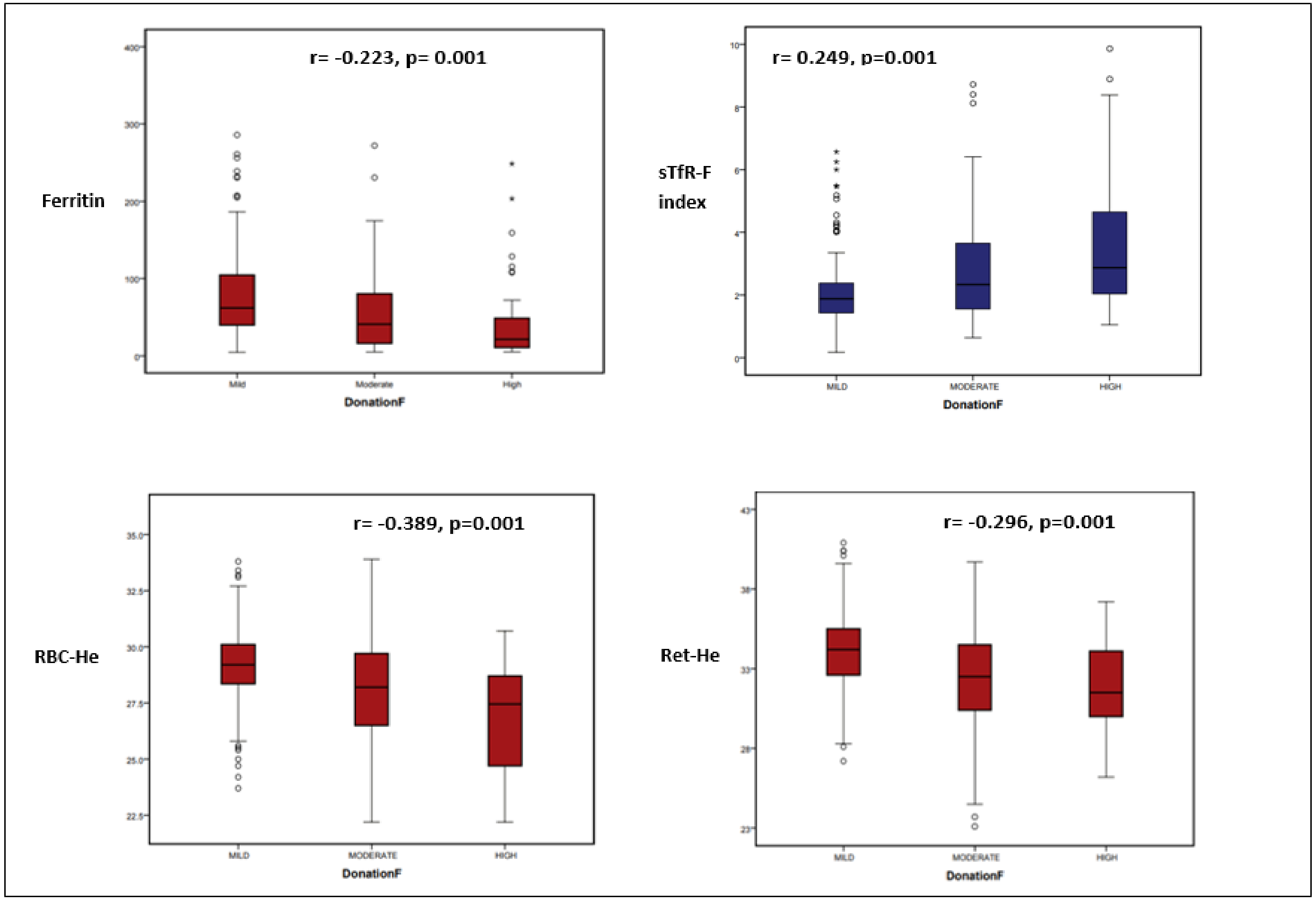

**Supplemental Figure 3- Correlation between platelet counts and MPV in iron-deficient donors (n=51)**

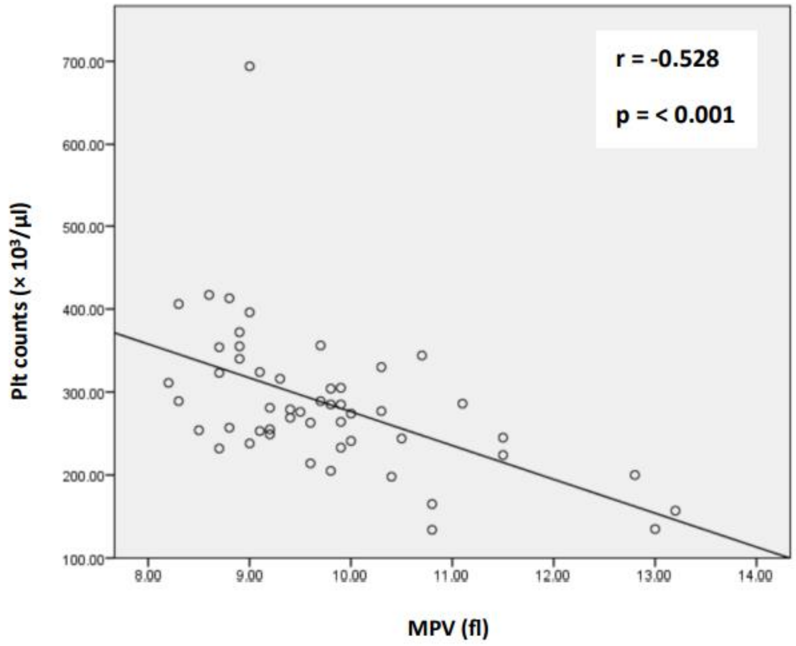
